# Alzheimer’s Disease Knowledge Graph Enhances Knowledge Discovery and Disease Prediction

**DOI:** 10.1101/2024.07.03.601339

**Authors:** Yue Yang, Kaixian Yu, Shan Gao, Sheng Yu, Di Xiong, Chuanyang Qin, Huiyuan Chen, Jiarui Tang, Niansheng Tang, Hongtu Zhu

## Abstract

**Background:** Alzheimer’s disease (AD), a progressive neurodegenerative disorder, continues to increase in prevalence without any effective treatments to date. In this context, knowledge graphs (KGs) have emerged as a pivotal tool in biomedical research, offering new perspectives on drug repurposing and biomarker discovery by analyzing intricate network structures. Our study seeks to build an AD-specific knowledge graph, highlighting interactions among AD, genes, variants, chemicals, drugs, and other diseases. The goal is to shed light on existing treatments, potential targets, and diagnostic methods for AD, thereby aiding in drug repurposing and the identification of biomarkers.

**Results:** We annotated 800 PubMed abstracts and leveraged GPT-4 for text augmentation to enrich our training data for named entity recognition (NER) and relation classification. A comprehensive data mining model, integrating NER and relationship classification, was trained on the annotated corpus. This model was subsequently applied to extract relation triplets from unannotated abstracts. To enhance entity linking, we utilized a suite of reference biomedical databases and refine the linking accuracy through abbreviation resolution. As a result, we successfully identified 3,199,276 entity mentions and 633,733 triplets, elucidating connections between 5,000 unique entities. These connections were pivotal in constructing a comprehensive Alzheimer’s Disease Knowledge Graph (ADKG). We also integrated the ADKG constructed after entity linking with other biomedical databases. The ADKG served as a training ground for Knowledge Graph Embedding models with the high-ranking predicted triplets supported by evidence, underscoring the utility of ADKG in generating testable scientific hypotheses. Further application of ADKG in predictive modeling using the UK Biobank data revealed models based on ADKG outperforming others, as evidenced by higher values in the areas under the receiver operating characteristic (ROC) curves.

**Conclusion:** The ADKG is a valuable resource for generating hypotheses and enhancing predictive models, highlighting its potential to advance AD’s disease research and treatment strategies.

## Background

Alzheimer’s disease (AD) is a neurodegenerative disorder characterized by progressive cognitive decline, memory impairment, and functional disability [1]. With an aging population, the prevalence of AD has been steadily rising, posing significant challenges to healthcare systems worldwide [2]. Alzheimer’s disease (AD) research has evolved significantly, expanding beyond the amyloid hypothesis to encompass tau pathology, neuroinflammation, and vascular factors [3–5]. Diagnostic advances include promising blood-based biomarkers and advanced neuroimaging techniques, while artificial intelligence enhances early detection [6–8]. Treatment approaches have expanded, including the controversial FDA approval of Aducanumab in 2021, alongside continued development of various anti-amyloid and tau-targeting therapies in clinical trials [9,10]. Prevention efforts focus on lifestyle interventions and vascular health, with a shift towards personalized medicine and recognition of AD subtypes [11–14]. Clinical trials also target earlier disease stages with novel designs to increase efficiency [15,16], while improved patient care through digital technologies and better management of behavioral symptoms complement biomedical research [17,18].

As the field of AD research rapidly evolves, it becomes increasingly crucial to synthesize and summarize information from the multitude of studies and published papers. This comprehensive approach allows researchers, clinicians, and policymakers to gain a holistic understanding of the current state of AD research and treatment. By consolidating findings from diverse areas such as disease mechanisms, diagnostic tools, treatment strategies, and care approaches, we can identify emerging trends, highlight promising avenues for future research, and inform evidence-based practices in AD management. Furthermore, regular summaries of the expanding body of knowledge facilitate the translation of research findings into clinical practice and policy decisions, ultimately advancing our collective efforts to combat this devastating disease.

One promising data mining method involves creating interaction triplets, consisting of three components: head entity, tail entity, and their relationship [19]. For example, let’s consider a sentence "PPARgamma may be a potential target for AD", we can obtain a triplet whose head entity is PPARgamma, the tail entity is AD, and their relationship is "potential target for". These triplets efficiently organize and make accessible the extensive knowledge embedded in the AD- related literature. By aggregating and examining these triplets, researchers can achieve a holistic view of AD research progress, paving the way for the construction of knowledge graphs that further illuminate the disease’s complexities.

In the biomedical domain, knowledge graphs are constructed through meticulous manual curation, seamless integration of existing databases, and innovative data-driven approaches. Many knowledge graphs, like Gene Ontology [20], Drug Bank [21], and UMLS [22], have been built through intense expert-led curation efforts. In addition, some knowledge graphs amalgamate various established databases, including DisGeNet [23], Hetionet [24], BioGrakn [25], and DemKG [26], to create comprehensive resources.

Specific to AD, there has been ongoing effort to develop AD-specific knowledge graphs. AlzPathway [27,28] is a notable example, offering a detailed pathway map of AD-related signaling pathways, curated from over a hundred review articles. The Alzheimer’s Disease Ontology (ADO) [29] stands out as the pioneering structured framework to systematize AD-related information, developed in line with the ontology building life cycle. Further contributing to the structured representation AD knowledge are the Alzheimer’s Disease Map Ontology (ADMO) [30], derived from AlzPathway, and the Alzheimer’s Disease Integrated Ontology (ADIO) [31], which merges ADO and ADMO. In addition to ontology development, efforts have been made to integrate multi-omics and heterogeneous biological networks for Alzheimer’s drug discovery. For instance, the Alzheimer’s Cell Atlas (TACA) [32] compiles transcriptomic data from over 1.1 million cells/nuclei across major brain regions and cell types, and integrates differential expression comparisons, protein-protein interaction modules, functional enrichment analyses, drug screening profiles, and cell-cell interaction analyses into an interactive web portal. AlzGPS [33] integrates multi-omics data and clinical databases for AD, offering curated multi-omics datasets, endophenotype disease modules, treatment information from FDA-approved drugs, literature references, clinical trial data, and interactive visualization tools to accelerate therapeutic development.

In recent years, there’s been a shift towards leveraging data mining techniques to extract AD insights from academic literature. Zhu [34]’s work exemplifies this trend by creating disease-specific knowledge graphs, including for AD, from PubMed abstracts, employing advanced models like Att-BiLSTM-CRF [35] for named entity recognition and a combination of BiLSTM . [36] and ResNet [37] for relation extraction. Similarly, Nian [38]’s research utilizes literature-derived knowledge graphs, extracting AD-related triplets from SemMedDB [39,40] to explore connections between AD and various entities, showcasing the growing emphasis on data-driven methodologies in constructing knowledge graphs for AD research.

The AD knowledge graph holds significant promise for advancing biomedical discoveries. Zhu’s creation of SDKG-11 [34], which encompasses knowledge graphs for five cancers and six non-cancer diseases including AD, showcases the application of diverse data processing methods. This work not only enhances existing models with multimodal reasoning but also proves its efficacy and broad applicability in uncovering new biomedical insights, especially in the realms of drug-gene, gene-disease, and disease-drug connections. Furthermore, Bang’s introduction of the DREAMwalk [41] framework marks a significant stride towards computational drug repurposing. By mapping drugs and diseases within a unified embedding space, DREAMwalk boosts the prediction of drug-disease associations, showing considerable promise for Alzheimer’s disease repurposing efforts.

Extracting information from text [42] is a pivotal step in knowledge acquisition, achievable through either open or supervised extraction methods. Open information extraction [43] autonomously discerns patterns in sentences without pre-supplied training data, employing tools like ReVerb [44], OLLIE [45], Stanford OpenIE [46], ClausIE [47], and SemRep [40]. In contrast, supervised information extraction [48] hinges on annotated datasets to derive semantic triplets, detailing entities and their interrelations. This approach typically involves named entity recognition (NER) and relation classification, executed either in stages or via integrated models to reduce inaccuracies. Recent advancements with models like SCIIE[49] and SpERT[50] have notably enhanced the accuracy and efficiency of supervised information extraction, further fueling the development and refinement of AD knowledge graphs.

Biomedical knowledge bases and the process of entity linking play crucial roles in structuring the vast amount of information extracted into coherent formats such as knowledge graphs. In these graphs, entities derived from triplets are connected to specific databases to clarify and resolve any ambiguities. Entity linking, also known as named entity disambiguation, can be conducted through supervised learning methods that utilize labeled identifiers, or through unsupervised techniques like string matching that associate textual mentions with unique database entities. Tools like TaggerOne [51] and Wikipedia2vec [52] leverage training data or embedding vectors to facilitate this linking process, harnessing the power of machine learning to enhance accuracy and relevance. Meanwhile, solutions such as QuickUMLS [53] and SciSpaCy [54] employ string matching algorithms, offering a direct approach to associate text with entities in UMLS or other biomedical databases. This method does not require training data, making it an accessible option for linking entities in a straightforward and efficient manner.

The goal of this paper is to develop a novel data mining method to extract information on AD from literature, primarily biomedical entities related to AD, and their relationships, to construct an AD knowledge graph. We employ a supervised strategy, harnessing crowdsourced annotations and GPT-4 [55] for textual enrichment to curate a training set for the SpERT[50] model training. We further delve into the knowledge graph’s attributes and its utility in hypothesis generation and disease prediction.

The key contributions of our paper are as follows:

1. Provide a brand-new human-annotated benchmark dataset for named entity recognition and relation classification specified in AD literature.
2. Propose a pipeline to construct a domain-specific ADKG with entities linked to reference databases.
3. Demonstrate the ADKG’s potential in predicting novel relationships and in forecasting AD’s disease prediction.

### Construction and content

We will first introduce the primary steps to construct the Alzheimer’s Disease Entity Relation Corpus (ADERC). Then, we will discuss on how we utilize the dataset to train an information extraction model for triplets’ extraction. Finally, we describe the approaches utilized to construct ADKG. The overall procedure of constructing ADKG is illustrated in Figure 1.

**Figure 1.**
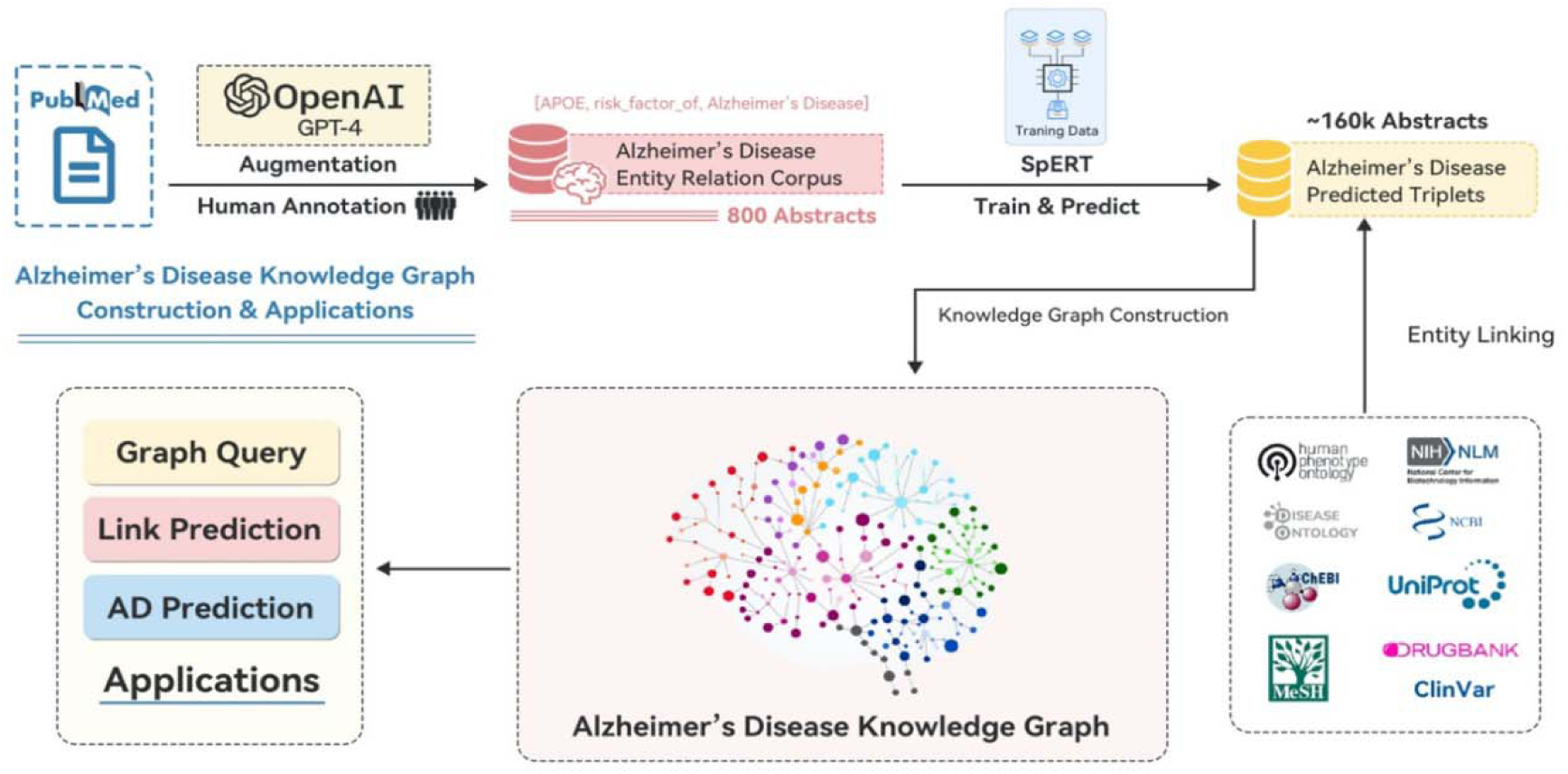
General Pipeline: corpus generation, model building, entity linking, ADKG construction, and applications

### Corpus Generation

A new dataset for AD-related information extraction, ADERC, is constructed from 800 PubMed abstracts related to AD. The overall procedure for generating the dataset is shown in Figure 1. Three steps are taken: abstract retrieval, pre-labeling using BERN[56], and annotation using BRAT[57].

We extracted 169,630 abstracts from PubMed (https://pubmed.ncbi.nlm.nih.gov/) utilizing the keyword ’Alzheimer’ through the Entrez function of the Bio package (as of 5/2/2021). The BERN tool [56], known for its prowess in biomedical entity recognition and normalization, was employed to initially tag named entities, assigning categories such as Gene/Protein, Disease, Drug/Chemical, Species, and Mutation, before the commencement of manual annotation.

Beyond the five entity types identified by BERN, annotators introduced an additional ’method’ category to classify methodological entities, exemplified by ’18F-FDG-PET’. An ’other’ category was also designated to encapsulate entities falling outside the predefined types. For the extraction of relationships, our focus was pinned on eight types: *treatment_for, treatment_target_for, help_diagnose, risk_factor_of, characteristic_for, hyponym_of, associated_with*, and *abbreviation_for*, which were selected to align with AD-related research inquiries. A meticulous manual review of 800 abstracts was conducted, leading to their comprehensive annotation via the BRAT tool [57]. This process resulted in a richly annotated corpus that encompassed both biomedical entities and their interrelations across 800 abstracts.

To bolster the training dataset for named entity recognition and relation classification, we utilized GPT-4 [55] to create textual variations via synonym substitution and phrase rephrasing, particularly targeting sentences that encapsulate relationships (refer to the supplementary section for detailed prompts used in generating this augmented data). Following rigorous manual reviews to ensure accuracy and mitigate biases, this enhanced dataset significantly contributed to elevating the model’s performance and minimizing the likelihood of overfitting.

### Information Triplet Extraction

We developed a unified model for simultaneous named entity recognition and relation extraction on ADERC, leveraging the SpERT [50] framework with SciBERT [58] embeddings. To refine our negative sampling approach, we introduced manually crafted negative instances. Beyond the standard practice of using texts without named entities or unrelated entity pairs as negative examples, we enriched the dataset by modifying positive instances through the substitution of entities and relations. This method is designed to achieve a balanced dataset, offering a comprehensive assortment of both positive and negative samples to reduce model bias and improve precision in detecting pertinent entities and their connections.

For model robustness, we partitioned the annotated corpus into training, validation, and testing sets. The model underwent training on the training set across a range of parameter configurations, including variations in relationship filtering thresholds, embedding dimensions, negative sample volumes, and dropout rates. The optimal parameter configuration was determined based on the best performance on the validation set. Employing this optimal setting, the finalized model was then applied to the entire corpus of unannotated data to systematically identify named entities and extract relation triplets.

### Abbreviation Resolution

In the development of knowledge graphs, handling abbreviations poses a notable challenge due to their potential for ambiguity, with the same abbreviation possibly representing different entities. For example, "ASD" could refer to "autism spectrum disorder" or "atrial septal defect." To mitigate such ambiguities, we’ve introduced a specific relationship type termed "*abbreviation_for*" in our annotation schema. This addition allows for the explicit representation of abbreviation relationships within the extracted triplets, significantly improving our ability to distinguish between entities. Our approach to resolving ambiguities involves first associating abbreviations with their full forms within the same abstract, thereby utilizing the broader textual context to aid in precise entity identification. The underlying principle is that the expanded form of an abbreviation offers a more comprehensive context crucial for accurate entity recognition and disambiguation.

### Entity Linking

The issue of inconsistent entity representations, whether across various abstracts or within different sentences of the same text, can introduce ambiguity. To address this, we have established a thorough entity linking procedure that coherently associates references to identical entities. This process utilizes an array of biomedical databases, each catering to specific types of entities. Our extensive database ensemble encompasses genes from NCBI Gene [59], proteins as outlined in UniProt [60], small molecules listed in ChEBI [61], pharmaceuticals detailed in DrugBank [62], phenotypes described in HPO [63], diseases cataloged in the Disease Ontology [64], mutations recorded in ClinVar [65,66], and other medical entities classified under MeSH. In this framework, every entity is allocated a unique identifier (ID) from its respective source database, enriched with detailed descriptions and additional information to support precise entity resolution and linkage. We employ simstring [67] for approximating string matching, comparing entity mentions in our extracted triplets against standard names and their synonyms in the reference databases, with matching scores serving as a measure of confidence.

### Knowledge Graph construction and the Confidence

A knowledge graph (KG), G(X, E) consists of nodes {X_l_, X_2_,…, X_N_ } E X and edges, {E_l_, £_2_,…, £_K_ } E E between nodes. In this study, to build a knowledge graph, G(X, E), for AD from the existing literature, we extract entities (nodes) and relationships (edges) from abstracts related to AD.

The ADKG is developed from the triplets extracted across all abstracts using the trained model. The knowledge graph’s construction involves two primary steps: creating nodes and establishing edges. In the node creation phase, abbreviations are resolved, and entities are identified through a process known as entity linking. Subsequently, for each identified triplet, edges are established, encapsulating the linked entities, the original PubMed ID, the spans of the entities within the text, and the matching scores. It’s common for multiple edges to exist between a pair of nodes. In the finalized ADKG, the nature of the directed edge connecting two nodes is determined by the predominant relationship types observed in the edges connecting the head and tail entities.

### Knowledge Fusion

Integrating external knowledge graphs with the Alzheimer’s Disease Knowledge Graph (ADKG) is essential to enhance the comprehensiveness and accuracy of the information available to researchers and practitioners [68]. This integration enriches the ADKG with diverse datasets, enabling more robust analyses and insights into Alzheimer’s disease. To preserve the integrity of the integrated knowledge graph, we included the sources of each entity and relationship. Additionally, we have developed a comprehensive mapping schema to facilitate the alignment of entities and relationships across different knowledge graphs. We integrated representative external databases like DisGeNET [23], The Human Phenotype Ontology (HPO) [63], DrugBank [62], PharmGKB (Pharmacogenomics Knowledgebase) [69], OMIM (Online Mendelian Inheritance in Man) [70], and STRING [71]. The full details of how we processed the external databases in the integration process is available in the supplementary materials.

Initially, a comprehensive manual examination of the external database’s schema and content is conducted to identify pertinent relationships for each of the external knowledge graph. Subsequently, entity linking techniques are utilized to map entities from the external database to their corresponding entities within the ADKG. Once entities are accurately linked, attention shifts to the selection of relationships. Specifically, relationships from the external database are selected if either the head entity (the origin of the relationship) or the tail entity (the destination of the relationship) exists within the ADKG. This selective approach ensures that only relevant relationships are integrated, thereby maintaining the integrity and relevance of the ADKG.

### Knowledge Graph Embedding

In developing the knowledge graph embedding model, we utilized various embedding techniques on the training set and determined the optimal parameters based on performance in the test set to create the final model for discovering new knowledge. The ADKG was divided into training (approximately 80%), validation (around 10%), and testing (near 10%) subsets. We began with a balanced distribution of relationships across these subsets, followed by manual adjustments to ensure all entities were represented in the training portion.

We evaluated several knowledge graph embedding (KGE) methods, including distance-based models like TransE [72], TransH [73], and TransR [74], the semantic matching-based ComplEx model [75], and the ConvKB model [76], which incorporates convolutional neural networks. Each model offers a unique approach to representing relationships and entities: (i) TransE treats relationships as translations in the embedding space, where the plausibility of a fact is the L1 or L2 distance between the sum of the head entity and relation vectors and the tail entity vector, embodying the concept of the relation ’translating’ the head to the tail entity. (ii) **TransH** projects entity embeddings onto relation-specific hyperplanes, accommodating entities’ differing roles across multiple relations. Its scoring function assesses fact plausibility based on the L1 or L2 norm after this projection, reflecting the translation principle on these hyperplanes. (iii) **TransR** separates entity and relation embeddings into distinct spaces, projecting entity embeddings into a relation-specific space before translation. The fact’s plausibility is measured by the L1 or L2 distance between the translated head entity and the tail entity within this relation space. **(iv) ComplEx** uses complex-valued embeddings to represent both symmetric and asymmetric relationships, with its scoring function being the real part of the dot product of complex embeddings, facilitating the modeling of diverse relation types, including hierarchical and reciprocal relations. (v) **ConvKB** employs a convolutional neural network over concatenated embeddings of head entities, relations, and tail entities to identify global interaction patterns. Its scoring involves convolution, a non-linear feature map, and a linear scoring layer, capturing intricate interaction patterns to predict new facts. By comparing the performances of these models, we selected the most effective one for facilitating knowledge discovery within the ADKG.

For all models, we conducted a comprehensive search for the optimal hyper-parameters, including choices for embedding dimensions (32, 64, 128, 256), learning rates (0.05, 0.005, 0.0005), batch sizes (128, 256, 512, 1024), and specific parameters for each loss function. For ConvKB, we additionally selected the number of filters from options 128, 256, and 512. We adopted the margin-based ranking loss function and implemented Bernoulli negative sampling [73] for training. The training process was capped at a maximum of 1000 epochs. During evaluation, we utilized a rank-based metric, specifically focusing on the mean rank of correct predictions, and applied a filtered setting [72] to account for known triples when ranking candidate triples. The primary criterion for model selection was the arithmetic mean rank, although we also reported Hits@10 as an additional measure to gauge model performance. Arithmetic Mean Rank is a measure used to evaluate the performance of models by calculating the average rank assigned by a model to a set of triplets. A lower arithmetic mean rank indicates better performance, as it suggests that the model is more accurately ranking items according to their relevance or importance. Hits@10 is a metric commonly used in information retrieval and recommendation systems to measure the proportion of relevant items that appear within the top-K results recommended by a model. In this case, Hits@10 specifically measures the percentage of relevant items that are included in the top 10 recommendations provided by the model. A higher Hits@10 score indicates better performance, as it suggests that more relevant items are being recommended within the top results.

## Results

### Statistics for ADERC and Model

The constructed dataset (ADERC) includes annotations for biomedical entities and their relations for 800 abstracts. These abstracts are retrieved from PubMed through query of "Alzheimer’s disease". The ADERC contains 20, 886 annotated mentions and 4, 935 relationships between these mentions. The original predictions on all the abstracts contains in total 3,199,276 entity mentions and 633,733 triplets, among which we identified 45,277 unique triplets between mapped entities after entity linking. Details of the types of entity and relationships are shown in Figure 2.

**Figure 2.**
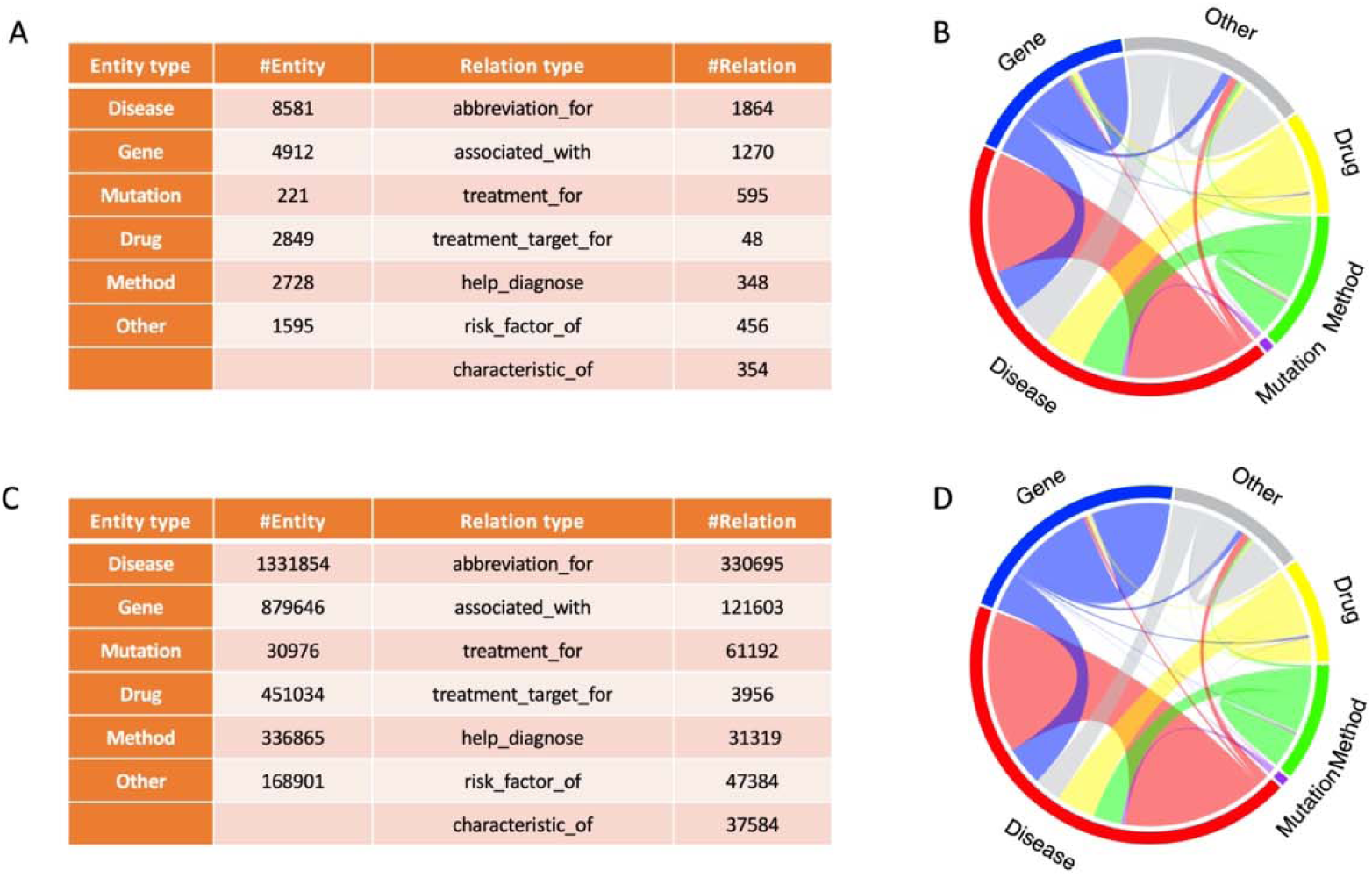
Comparative Visualization of Biomedical Entity and Relationship Distribution for ADERC (A and B) and ADKG (C and D). (Not all the tail entities are Alzheimer’s disease.)

**Figure 3.**
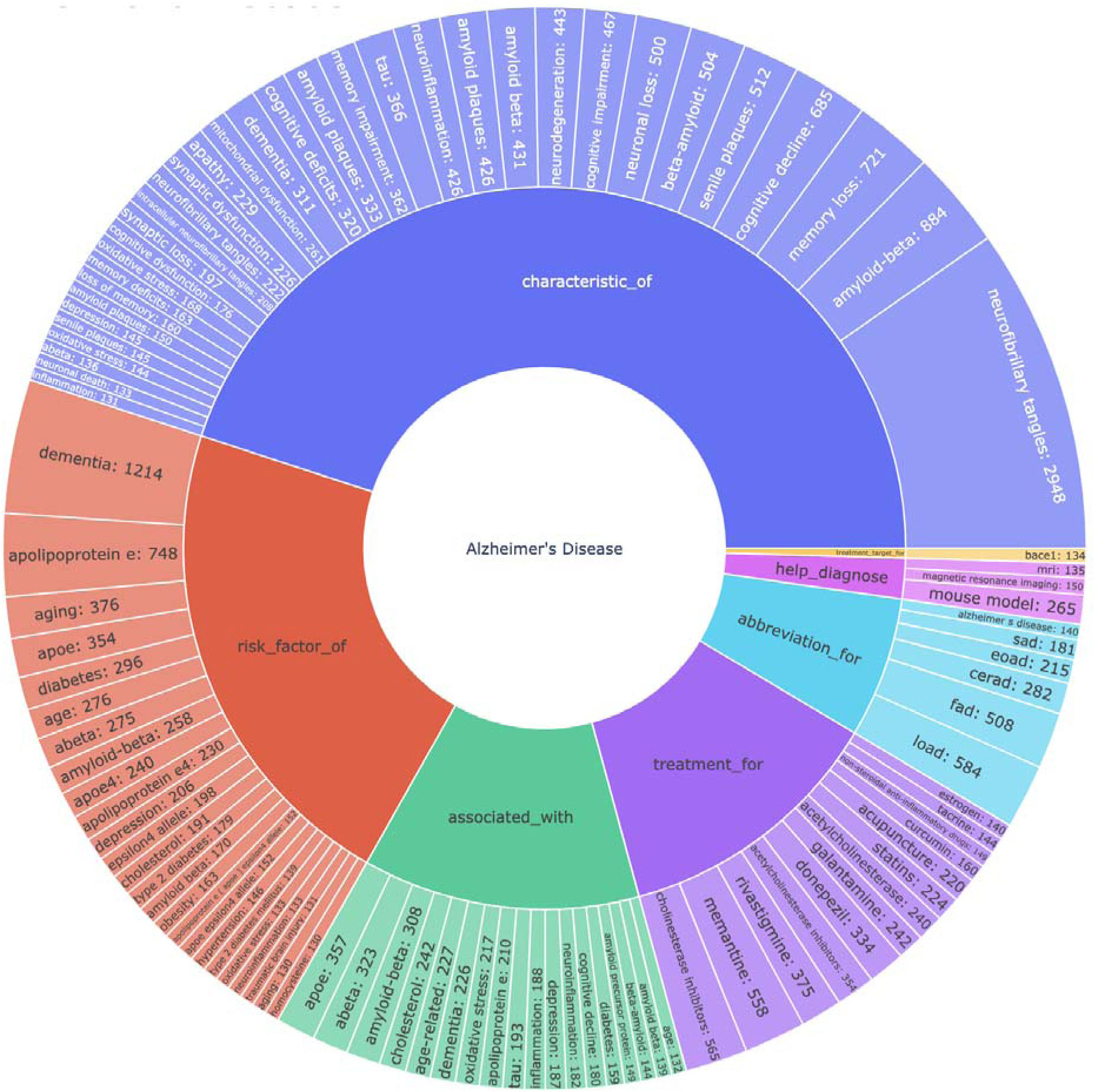
Top 100 relationships related to Alzheimer’s Disease with the degree the number of sentences predicted to have the relationship

### NER&RE Model Performances

We present a detailed comparison of the precision, recall, and F1 score metrics for three different models: Entity Recognition, Relation Extraction, and a Joint model that combines the two tasks. Precision and recall are reported in both micro and macro averages, offering insights into the models’ performance across individual instances (micro) as well as across different classes (macro). For Entity Recognition, the model demonstrates strong performance with a micro F1 score of 87.2% and a macro F1 score of 86.1%, indicating consistent accuracy across various entity types. Relation Extraction shows lower scores across all metrics, with a micro F1 score of 67.1% and a macro F1 of 66.3%, reflecting the increased challenge of this task. The Joint model, which tackles both entity recognition and relation extraction simultaneously, understandably has lower scores than the individual tasks with a micro F1 of 61.4% and a macro F1 of 61.5%, suggesting that combining tasks may introduce additional complexity.

### Precision of ADKG

Assessing the performance of a knowledge graph, such as the ADKG, necessitates a meticulous evaluation, particularly focusing on the precision of the predicted triplets to confirm the graph’s effective encapsulation of the sourced information. This evaluation involves selecting a random sample of 100 sentences from the relevant literature and examining the triplets that the ADKG derives from these sentences. Each triplet’s accuracy is scrutinized to verify its faithful representation of the sentence’s content. Precision is quantified as the proportion of accurate triplets in relation to the total number of triplets extracted. This rigorous validation process initially employs GPT-4 and is subsequently cross-verified by domain specialists. In our analysis, we found that 94 out of 137 triplets (68.61%) were correctly represented in the ADKG, demonstrating its efficacy in capturing pertinent information.

### ADKG Featured Important Relationships for AD

In leveraging the ADKG, we observe recurring patterns, such as the well-documented link between the APOE gene and AD as a significant risk factor. The ADKG facilitates the discovery of complex relationships between AD and various entities. For example, when investigating genetic factors associated with AD, a query for ’AD – genes’ reveals 5,932 interactions connecting AD to 1,030 genes/proteins. This collection includes genes identified by the ADSP Gene Verification Committee as having a potential impact on AD risk or offering protective effects [77]. In terms of pharmacological connections, our analysis brought to light 5,665 interactions between AD and 1,061 different drugs/chemicals. This comprehensive network highlights numerous avenues for potential therapeutic interventions and illustrates the value of the ADKG as a resource for generating and validating hypotheses within Alzheimer’s disease research.

Furthermore, the analysis of disease comorbidity within the ADKG highlights notable connections between AD and other medical conditions. We identified 5,130 interactions that associate AD with 248 different diseases. Notably, Alzheimer’s Disease (DOID: 10652) is strongly linked to conditions such as Mild Cognitive Impairment (DOID: 0081292), Diabetes Mellitus (DOID: 9351), and Obesity (DOID: 9970). These associations are crucial for understanding the co-occurrence of these diseases and can provide essential insights for developing clinical approaches to diagnose, treat, and manage AD alongside its comorbidities effectively.

### Results of Knowledge Graph Embedding

Utilizing a grid search across all potential parameters, we identified the optimal configurations for knowledge graph embedding, assessed via the mean rank metric for the selected KGE models on the test set. The experimental outcomes, as presented in Table 1, indicate that the ConvKB model outperforms others, achieving the most favorable mean rank results on the test set.

**Table 1.**
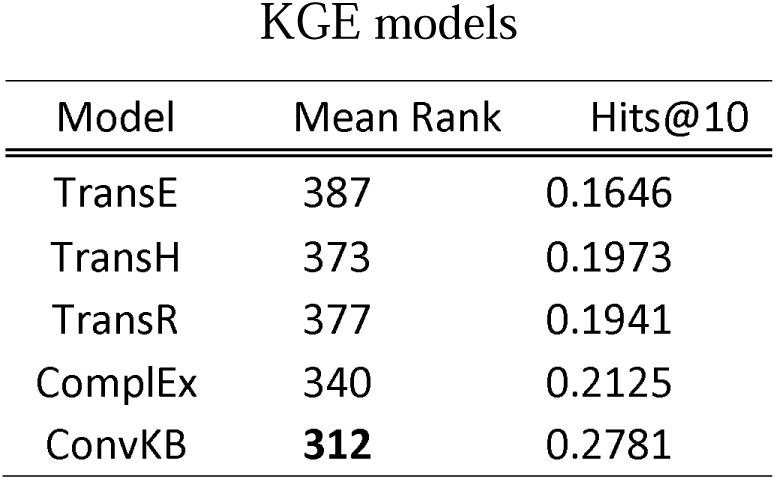
Knowledge Graph Embedding performance of the best setting on test set for different KGE models.

### Link prediction results reveal interesting findings

To uncover potential triplets not currently represented in the ADKG, we utilized the entire ADKG as a training dataset. Selecting ConvKB as the optimal model, we applied it with the most effective parameter configuration to train a KGE model on all ADKG triplets. We sought to discover new connections by calculating scores for all conceivable head-tail-entity triplets, ranking them based on these scores. This ranking of potential triplets offers insights into prospective or previously unidentified relationships among ADKG entities.

We compiled a list of new gene-disease relationships that emerged from link prediction, prioritizing top-ranking triplets that suggest associations between specific genes and diseases (Table 2). These diseases span a range of neurological and inflammatory conditions, including amyloidosis, neurodegeneration, and gastrointestinal inflammation, hinting at the implicated genes’ involvement in these disorders. Notably, *CHI3L1* is implicated in connections with neurodegeneration and hippocampal atrophy, a finding corroborated by recent publications not included in our initial PubMed dataset. This highlights the ADKG’s utility in revealing novel relationships and underscores the potential for advancing our understanding of complex diseases through knowledge graph analysis.

**Table 2.** Top inferred triplets inferred from ADKG using ConvKB (red PubMed evidence indicates that the source is not included in our corpus)

| Type | head | tail | rank | score | Pubmed Evidence |
| --- | --- | --- | --- | --- | --- |
| gene-disease | *APOE* | amyloidosis | 38 | 44.80191 | Many |
|  | *CHI3L1* | Neurodegeneration | 45 | 44.66664 | 35234337 |
|  | *MFN2* | Abnormality of mitochondrial metabolism | 46 | 44.66243 | 30649465 |
|  | *APOE* | tauopathy | 59 | 44.33801 | Many |
|  | *CRP* | Gastrointestinal inflammation | 64 | 44.21387 | Many |
|  | *HMOX1* | progressive supranuclear palsy | 74 | 43.9729 |  |
|  | *IL6* | major depressive disorder | 95 | 43.41697 | Many |
|  | *NEFL* | Neurodegeneration | 99 | 43.35546 | Many |
|  | *UBB* | neurodegenerative disease | 101 | 43.32749 | Many |
|  | *CHI3L1* | Hippocampal atrophy | 109 | 43.21873 | 35234337 |

### ADKG empowers Alzheimer’s Disease prediction using UK Biobank data

In this study, we sought to evaluate the predictive power of ADKG in identifying AD using the extensive data resources of the UK Biobank [78]. The UK Biobank offers an extensive biomedical database that includes genetic and health information from nearly half a million UK residents. Our methodology involved retrieving relevant data from the UK Biobank as of March 6, 2023, which included protein expression profiles at enrollment, lifestyle information, and detailed medical histories crucial for AD prediction. Our analysis focused on predictors such as protein abundance, environmental factors, and lifestyle variables, to predict the diagnosis of AD indicated by the case group and control group. The predictive variables encompass 1463 proteins, APOE4 variant, race, education level, social demographics, smoking status, physical activity level, diet, alcohol usage, sleep, and memory.

We identified UK Biobank participants diagnosed with AD as the case group based on hospital admission electronic health records (EHRs), using ICD-9 and ICD-10 codes from linked records or data from death registers. For our control group, we selected individuals without any dementia-related symptoms according to ICD-10 codes, specifically excluding codes with F00 (Dementia in Alzheimer’s disease), F01 (Vascular dementia), F02 (Dementia in other diseases classified elsewhere), F03 (Unspecified dementia), F05 (Delirium, not induced by alcohol and other psychoactive substances), G30 (Alzheimer’s disease), G31 (Other degenerative diseases of nervous system, not elsewhere classified), and G32 (Other degenerative disorders of nervous system in diseases classified elsewhere), and excluding individuals with any form of dementia reported by the UK Biobank. The complete dataset of 52164 cases and 541 controls was then randomly divided into training and validation (90%), and testing (10%) subsets for analysis.

We conducted a comparative evaluation of phenotypes associated with AD as indicated in ADKG versus those identified through conventional screening, particularly in their ability to predict AD. For phenotypes referenced from ADKG, we selected AD-related genes/proteins in ADKG and manually reviewed other entity types linked to AD (such as lifestyle variables and environmental factors) in the UK Biobank health records.

In developing the predictive model for AD, we utilized both logistic regression and XGBoost algorithms [79]. To prepare for model training, we tackled the issue of missing data in the normalized protein expression profiles by employing mean substitution [80], ensuring the completeness and reliability of the data essential for the logistic regression model, given the complexity of missing data patterns. The relatively low occurrence of AD in the population, leading to an imbalanced case-control ratio in our dataset, prompted us to use the ROSE package [81] for oversampling, achieving a more equitable distribution of cases and controls in the training dataset.

For the selection of variables without relying on domain-specific knowledge, we set p-value thresholds at various levels (0.05, 0.005, 0.0005, 0.00005, 0.000005, 0.0000005), each yielding a different set of predictors. Leveraging information from ADKG, we incorporated a subset of AD-associated genes and other pertinent variables such as age, the presence of the *APOE* ε*4* allele, and cognitive memory scores, resulting in a comprehensive dataset comprising 214 variables for the analysis.

The integration of domain-specific knowledge from the ADKG substantially improved the model’s predictive accuracy, as demonstrated by an increase in the Area under the Receiver Operating Characteristic (ROC) curve from 0.9025 to 0.9137 for the ADKG-enhanced model.

Moreover, the application of the XGBoost algorithm with ADKG-derived predictors achieved an even higher AUC of 0.928, underscoring the potency of sophisticated machine learning techniques in refining AD predictive models. These results highlight the pivotal role of domain-specific knowledge in augmenting model performance. This comprehensive evaluation process is depicted in Figure 4.

**Figure 4.**
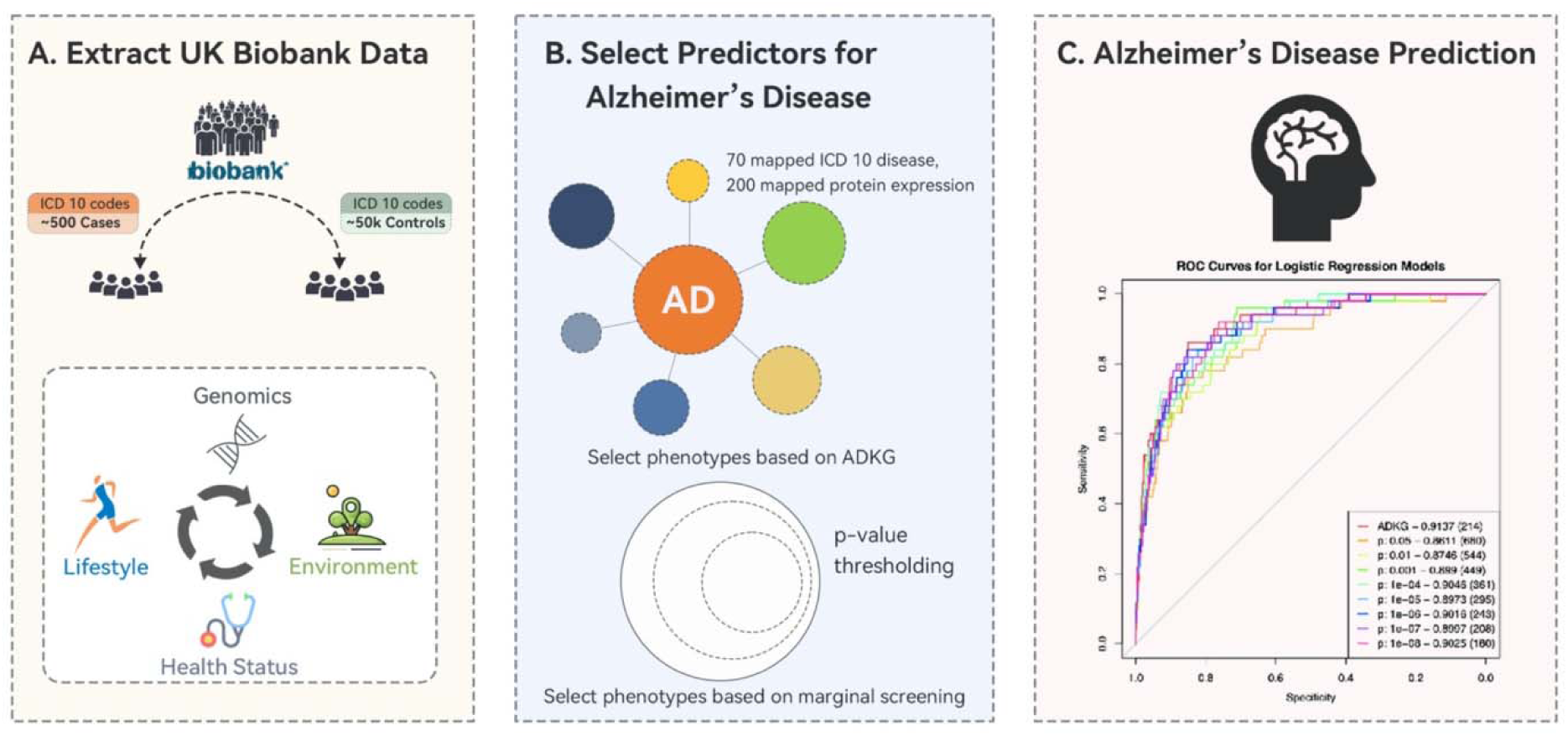
Efficacy of ADKG in AD prediction with UK Biobank Data

## Discussion

In this study, we unveil a cutting-edge data mining approach for developing the ADKG, derived from triplets extracted from academic abstracts. This knowledge graph serves as a pivotal tool for drug repurposing and identifying biomarkers pertinent to AD. Additionally, we introduce the ADERC, a uniquely human-annotated dataset tailored for research in AD knowledge graphs. Our methodology delves into the ADKG’s attributes, employing it for data retrieval and showcasing its utility in discovering new connections through link prediction techniques enabled by knowledge graph embedding methods. A cornerstone of our research is the practical use of the ADKG in the predictive modeling of AD, capitalizing on the comprehensive data available within the UK Biobank.

While our framework demonstrates considerable promise, there are avenues for enhancement. Beyond technical refinements, a critical area for expansion involves broadening the data sources beyond abstracts to include full-text articles, as well as data from supplementary materials like tables and notes, to enrich the depth of information extracted. Further granularity in classifying entity and relationship types could also provide deeper insights. For example, differentiating the ’gene’ category into more specific types such as gene, protein, and RNA could offer more precise understanding. Recognizing negative relationships is equally important, as it can help identify erroneous conclusions in AD research. Such advancements would necessitate increased annotation efforts to generate enough annotated training samples for each detailed category, ensuring the model’s ability to accurately discern these patterns.

Regarding the application of the ADKG, we have outlined various potential uses and presented case studies to illustrate how the ADKG can enhance traditional tasks. It’s important to note, however, that the scope for improving traditional tasks with knowledge graph insights extends beyond the examples provided in our manuscript. The emergence of large language models opens up even more possibilities. One notable application could involve integrating ADKG data into a question-and-answer (Q&A) engine powered by large language models, thereby making Alzheimer’s Disease information more readily accessible to the general public.

### Key Points

1. The study focuses on developing an Alzheimer’s Disease Knowledge Graph (ADKG) by extracting relationships between genes, variants, chemicals, drugs, and diseases related to Alzheimer’s from 800 PubMed abstracts, using GPT-4 for text augmentation.
2. A joint model that integrates named entity recognition (NER) and relationship classification was trained and used to parse unannotated abstracts. Reference biomedical databases were used for entity linking, enhanced by abbreviation resolution techniques.
3. The ADKG enabled Knowledge Graph Embedding models to generate high-quality, evidence-supported hypotheses. The predictive models using ADKG, when tested on the UK Biobank data, showed superior performance with higher areas under the ROC curves compared to other models.

### Availability of data and material

The datasets generated during and/or analyzed during the current study are available in the Zenodo repository https://doi.org/10.5281/zenodo.5770100. The knowledge graph ADKG is accessible via our developed website https://biomedkg.com/ for easy query and visualization.

## Authors’ contributions

YY, KY and HZ designed the analysis plan and interpreted the analysis results. YY carried out the analysis and draft the manuscript. SY reviewed the process and proposed improvement. SG, DX, CQ, HC, JT, and NT annotated the dataset.

All authors read and approved the final manuscript.

