## Supplementary Materials for "Alzheimer’s Disease Knowledge Graph Enhances Knowledge Discovery and Disease Prediction"

**Supplementary Note**

### GPT augmentation

We used GPT for text augmentation to increase training samples for named entity recognition (NER) and relation classification by selecting a portion of the generated data and feeding it to the model as input. GPT generated new variations of the input text using a variety of techniques, such as synonym replacement, random insertion, and text masking, among others. Since relabeling for the whole augmented abstract is a demanding task and relations are more in need of augmentation, we only augment the sentences that have at least one relation. Furthermore, we manually evaluated the quality of the augmented data and ensured that it doesn't introduce any biases or errors. We then use these augmented labeled sentences as additional training data for our joint NER and relation extraction models. By increasing the diversity and quantity of our training data, we improved the performance of our models and reduced the risk of overfitting.

**Example input given to GPT:**

Rewrite the sentence and keep the listed strings unchanged. The sentence is: 18F-FDG PET Improves Diagnosis in Patients with Focal-Onset Dementias. The strings that do not change are: 18F-FDG PET, Focal-Onset Dementias. Return the rewritten sentences.

### Annotation guideline

With the challenges met in annotation trials, we carry out the following annotation guideline after careful consideration. Some of the practice follows the annotation guidelines from previous works [1,2]. Annotator is supposed to revise BERN results by adding and correcting tags with spans if entities are missing labels or labeled wrongly, and then annotate relations between tagged named entities within a sentence.

Table 3 Entity Types and Relation types (A, relation, B)

| Entity Type | Description | Example |
| --- | --- | --- |
| Gene/Protein | Gene or protein | Abeta40, Abeta42, beta-amyloid, … |
| Disease/Symptom | Disease or symptom | Alzheimer's disease, … |
| Drug/Chemical | Drug or chemical | memantine, ChEIs, … |
| Mutation | Genetic mutation | rs4291, rs4309 |
| Method | Tools, models, protocols, assessment tools, treatment therapies, … | (18)F-FDG PET imaging, Transgenic mouse model, routine pharmacotherapy, … |
| Other | Other related terms | tau hyperphosphorylation, Abeta deposition, … |
| Relation Type | Description | Example |
| Treatment_for | A is a (potential) treatment for B; A could be used to treat B | The use of guanfacine (Intuniv) for prefrontal cortical disorders… |
| Treatment_target_for | A is (potential) treatment target for B | PPARgamma may be a potential target for AD treatment… |
| Help_diagnose | A can help diagnose B | 18F-FDG PET Improves Diagnosis in Patients with Focal-Onset Dementias… |
| Risk_factor_of | A is a risk factor of B; A causes B | Ageing of the brain is the major risk factor for neurodegenerative disorders… |
| Characteristic_for | B is characterized by A; A is (pathological) hallmarks of B | … iNPH) is a neurological disorder characterized by gait disturbance … |
| Hyponym_of | A is a hyponym of B, B is a type of A | … neurodegenerative disorders such as AD… |
| Abbreviation_for | A is an abbreviation for B | Dementia with Lewy bodies (DLB) is … |
| Associated_with | A is associated with B | Our study further confirmed the association of GAB2 and AD… |

**Guidelines to Tag Entities**

1. Nested spans:
   a. longer spans belong to pre-defined categories is preferred
   b.
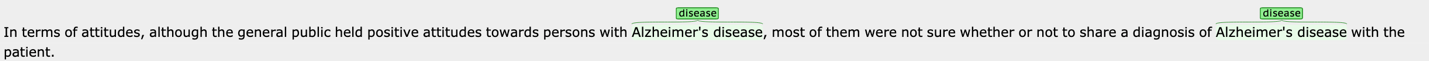
A sub-span within a longer annotated entity is tagged only when it has relationships with other entities outside the sub-span
2.
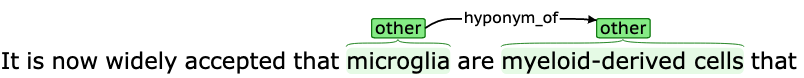
'Other' type is only tagged on entities when they have relationships with other entities.
3.
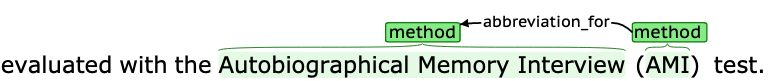
Determinators (e.g., a, an, …) or adjective pronouns (e.g., this, its, these, such, …) are all excluded from annotation
4. Adjective before a potential entity (a widely used term):
   a.
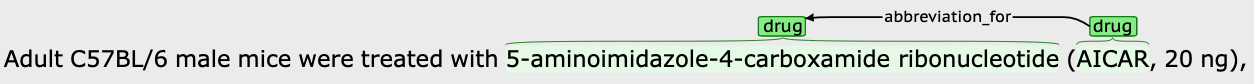
if the adjective is widely used before the entity or changes the meaning of relations tagged with it, include the adjective in an entity.
5. Noun after a potential entity (a widely used term):
   a. If including the noun into the potential entity changes its meaning or fulfil a relation tagged with it, include the noun.
   b.
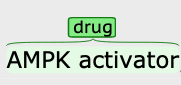
If including the noun does not change its meaning or fulfil a relation tagged with it, do not include the noun.

### Guidelines to Tag Relations

6.
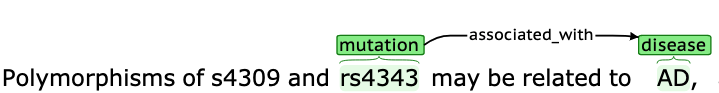
When trying to tag a relation into 'associated_with' category, only tag it if there are keywords in the following set: {"associated", "related", "correlated", "intetrwined", "interrelated"}.
7. Do not tag a relation when the relation is stated in doubt, in a hypothesis, or in a negative or questioning sense.

### Integrated External Knowledge Graphs

### Integrating various external knowledge graphs with the Alzheimer's Disease Knowledge Graph (ADKG) significantly enhances the richness and utility of the information available for Alzheimer's disease research. This integration requires careful selection of relevant knowledge graphs, ensuring compatibility in terms of entity types and relationship types.

### 1. DisGeNET [3]: DisGeNET is a discovery platform that integrates information on gene-disease associations from various sources, providing a valuable resource for the study of human diseases. It includes entity types such as Variants, Gene/Protein and Disease, and relationship types like associated_with. The relationships are retrieved by the R package disgent2r [4], which provides gene query for associated diseases, disease query for associated genes and variants, and variant query for associated diseases.

### 2. DrugBank [5]: DrugBank is a unique bioinformatics and cheminformatics resource that combines detailed drug data with comprehensive drug target information. The entity types in DrugBank include Drug/Chemical and Gene/Protein. These gene/protein entities are associated with Drug/Chemical entities as carriers, enzymes, transporters, and targets. These associations provide crucial insights into the interactions between drugs and proteins, facilitating a deeper understanding of drug mechanisms, metabolic pathways, and therapeutic applications. To represent all these associations in a standardized manner, the term "associated_with" is used.

### 3. The Human Phenotype Ontology (HPO) [6]: HPO provides a standardized vocabulary of phenotypic abnormalities encountered in human disease, facilitating the exchange of phenotypic data. HPO includes entity types such as Disease, Gene/Protein, and phenotypes. Its relationship gene-disease relationships are mendelian, polygenic, and unknown, and therefore we interpreted the first two types as risk_factor_of, and the last type associated_with. Other relationships are not labeled, and therefore we recorded them as associated_with.

### 4. PharmGKB (Pharmacogenomics Knowledgebase) [7]: PharmGKB curates knowledge about the impact of human genetic variation on drug responses, providing information on gene-drug-disease relationships. PharmGKB includes entity types such as Gene/Protein, Drug/Chemical, and Disease, and relationship types as associated_with.

### 5. OMIM (Online Mendelian Inheritance in Man) [8]: OMIM is a comprehensive, authoritative compendium of human genes and genetic phenotypes that is freely available and updated daily. OMIM includes entity types such as Gene/Protein, Disease, and Mutation, with relationship types we inherited as associated_with.

### 6. STRING (Search Tool for the Retrieval of Interacting Genes/Proteins) [9]: STRING is a database of known and predicted protein-protein interactions, encompassing both direct (physical) and indirect (functional) associations. It integrates data from various sources, including experimental repositories, computational prediction methods, and public text collections, providing a comprehensive view of protein networks that supports functional discovery in genome-wide experimental datasets. To ensure high confidence in the data, we restrict our downloads to interactions involving "Homo sapiens" and select only those associations with a combined score greater than 0.7 and interpret them into “associated_with” relationships. The combined score in STRING is computed by aggregating evidence from various types of interaction data, including neighborhood (genomic context), fusion (gene fusions in certain species), co-occurrence (genes found together across organisms), co-expression (correlated expression levels), experimental (biochemical evidence), database (curated information), and text mining (co-mentions in literature). This scoring system ensures that reported interactions are robust and reliable, making STRING a valuable resource for understanding protein functions and interactions in human biology.
